# An optimal set of inhibitors for Reverse Engineering via Kinase Regularization

**DOI:** 10.1101/2020.09.26.312348

**Authors:** Scott Rata, Jonathan Scott Gruver, Natalia Trikoz, Alexander Lukyanov, Janelle Vultaggio, Michele Ceribelli, Craig Thomas, Taran Singh Gujral, Marc W. Kirschner, Leonid Peshkin

**Affiliations:** Department of Systems Biology, Harvard Medical School, Boston, MA 02115, USA; Division of Human Biology, Fred Hutchinson Cancer Research Center, 1100 Fairview Ave N, Seattle, WA 98109, USA; Department of Pharmacology, University of Washington, Seattle, WA 98195, USA; Division of Preclinical Innovation, NCATS/NIH, Rockville, MD 20850, USA

## Abstract

We present a comprehensive resource of 257 kinase inhibitor profiles against 365 human protein kinases using gold-standard kinase activity assays. We show the utility of this dataset with an improved version of Kinome Regularization (KiR) to deconvolve protein kinases involved in a cellular phenotype. We assayed protein kinase inhibitors against more than 70% of the human protein kinome and chose an optimal subset of 58 inhibitors to assay at ten doses across four orders of magnitude. We demonstrate the effectiveness of KiR to identify key kinases by using a quantitative cell migration assay and updated machine learning methods. This approach can be widely applied to biological problems for which a quantitative phenotype can be measured and which can be perturbed with our set of kinase inhibitors.

## Introduction

Our knowledge of molecular circuitry traditionally grows one element at a time by a history of previous successes and failures that can hardly be characterized as unbiased. The promise of reverse engineering in systems biology is unbiased discovery of genes and pathways most causally related to the phenotypes we wish to explain. By unbiased we mean that it is not overly influenced by chance observations, the methods available for discovery or the reagents at hand. One example of the power of such unbiased large-scale approaches is the mutant screen, such as the ones that opened up so many avenues of discovery using chemical mutagenesis in Drosophila and analyzed as developmental defects in the larvae (Nüsslein-Volhard and Wieschaus 1980). Much of the functional knowledge of fly and general metazoan developmental biology began in this random unbiased way (though the analysis of these mutant phenotypes was hardly random). These genetic approaches were spectacularly successful but for other phenotypes a pharmacological approach would have its own advantages: post-transcriptional events such as protein translation, modification, and degradation can be explored; quantitative phenotypes can be assessed in a dose-dependent manner; inhibitors are generally portable to different cell types and organisms; and it is possible that discovery could lead directly to useful drugs.

Drugs have several imitations. Many drugs are optimized for a “one drug, one target” approach. The rationale is that disease is caused by mutation affecting the activity of one enzyme; if only that enzyme could be inhibited the disease could be managed. Not only is this view of disease misguided but the idea of a drug having only one target is as well. No matter the potency and specificity of a drug it will inevitably have ‘off targets’; e.g. what is labelled as a cyclin-dependent (CDK) 1 inhibitor will generally also inhibit other members of the CDKs amongst others. Not only is the one drug one target not as reliable as “one gene, one enzyme” but finding the target of a drug is not as automatic as finding the genetic target of a mutation has become. In fact this is the limitation of pharmacologic screening; you not only have to find one target, which is hard enough, but you have to find and understand all the other ones. Yet the experimental advantages of pharmacology are worth trying to bridge the drug/target divide. Furthermore since lack of specificity is usually not random (such as all CDK enzymes) there are phenotypes you can pick up that could not easily be seen if drugs were individually targeted to a single gene product. This has been the experience of gene knockouts in animals like the mouse. In the early days genes that were identified by spectacular overexpression phenotypes in mammalian cells like myoD, disappointingly had minor or irrelevant phenotypes in the knockout mouse. For this reason we sought a more quantitative and less qualitative way to exploit pharmacology for gene identification in a specific process. We turned this problem (quantitative not qualitative, polyspecific not specific) into a rationale for a new approach to exploit polypharmacology to deconvolve protein kinases involved in cell biological phenotypes (Taranjit Singh Gujral, Peshkin, and Kirschner 2014). We overcame the ‘problem’ of the lack of kinase inhibitor specificity with the realization that if we can quantitatively determine what the off targets are, then we can use that information to identify the kinases involved in a particular cellular phenotype. The outcome would not be a single gene but rather a rank-ordered list of genes participating in that phenotype. In that way the outcome can convey a great deal of information about the circuits controlling that phenotype. This idea is depicted in Figure 1, where drugs inhibit kinases and a quantitative cellular phenotype to differing degrees. The phenotype is then regressed against all kinases and by finding the right kinase, or more likely the right rank-ordered list of kinases, that contribute differentially to the formation of that phenotype, important features of how the phenotype is generated are revealed. Kinases are particularly appealing because of the large number of drugs available to target them and because they participate in practically every cell biological process. Their centrality is not unrelated to the medical importance they play in so many pathways. We should note that although we chose to look at kinase inhibitors, this method could apply equally well for other collections of inhibitors.

**Figure 1:**
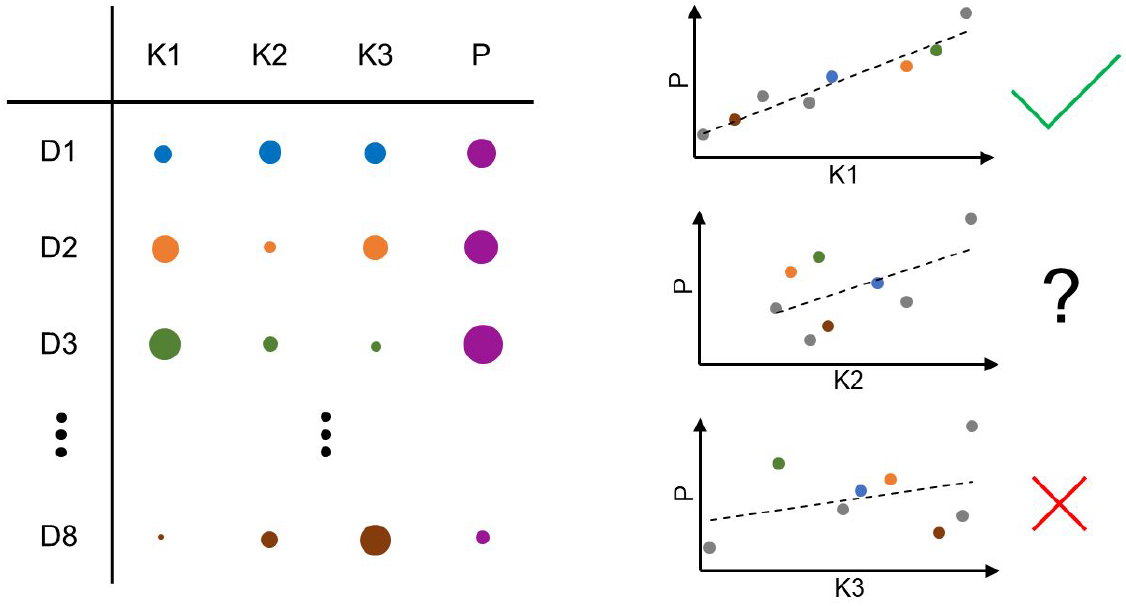
Concept of kinome regularization. Toy model of kinome regularization. Drugs D1–D8 inhibit kinases K1–K3 to a varying degree and affect the cellular phenotype, P. Circle sizes depict residual kinase activity and quantitative phenotypic values. By plotting P against the activity of each kinase in turn, where each point on the plot corresponds to a single drug, we can regress P against each kinase and determine regression coefficients. From these simple plots we can observe that our optimal drug set must be capable of inhibiting the kinases to varying degrees to run a successful regression analysis.

We previously applied this “kinome regularization” (KiR) using a publicly available tabulation of kinase inhibitor information (Anastassiadis et al. 2011) and a quantitative cell migration assay (Taranjit Singh Gujral, Peshkin, and Kirschner 2014). The original map of kinase inhibitor specificity was limited in many ways, so we set out to improve our optimal drug set. In principle there should be a minimal set of drugs whose multiple specificities could be used efficiently to explore the entire kinase space, so that one could convert the phenotypic responses to this smaller set of drugs into a rank ordered set of possible kinase targets. With this information we could identify the kinases and the underlying molecular mechanism responsibles for a given phenotype, using a minimal number of experiments. For this we need to know the quantitative relationship between each inhibitor and each kinase in the genome for a large number of drugs so we can select a minimal representative subset. We have data on 71 % of protein kinases. We have a relatively large amount of information on inhibitor specificity. As kinase inhibitors were developed, medicinal chemists often measured the relationship between chemical structure and inhibitor potency. These efforts were often focused on characterization of the protein target and occasionally a small number of relevant off-target kinases. Quantitative, high-throughput characterization of kinase inhibitors began in 2005 by Fabian et al. (Fabian et al. 2005) who used binding assays.(In simple kinetic models inhibition constants are genuine binding constants; they are not like Km, a ratio of rate constants). Since then numerous formats have been introduced including enzymatic determination of inhibition and measurements of inhibitor affinity. It has now become routine to characterize inhibitors against large portions of the protein kinome and these efforts have produced large sets of data on a wide range of both kinases and inhibitors (Gao et al. 2013; Davis et al. 2011; Elkins et al. 2016; D. H. Drewry et al. 2017). The focus of these studies largely remains confined to characterization of off-target effects in drug discovery and attempts to systematically identify the factors that govern kinase susceptibility to inhibition. They are part of a large pharmaceutical effort that, at least originally, aimed to identify very specific inhibitors, which has been of limited success. Despite the large volume of kinase-inhibitor data available, a key remaining limitation for systems level insight and machine learning approaches is the incompleteness of the data, where no single kinase is tested against every inhibitor nor is any particular inhibitor tested against every kinase. Furthermore, not all assays are run under the same conditions in a consistent, uniform manner. Such incompleteness and inconsistency can lead to bias in resulting analyses. The result is that to use this information effectively we had to repeat the very large kinase by inhibitor affinity (inhibition constant) behavior under uniform conditions. In our final KiR 3 data reported here this involved more than 25,000 assays each done at 10 concentrations, some repeated if there was noise in the measurement or other problems; in short, a significant amount of work.

Since our first application of KiR at least four more sets of kinase inhibitors have been published for human protein kinases. There are two main reasons we went ahead and profiled more inhibitors: more coverage in our set (369 kinases compared with 234 by Gao et al.; 257 compounds compared with 72 by Davis et al.; multiple doses compared with single doses) and, perhaps more importantly, a completely different motivation for building the set. Drewy et al. sought to seed collaborations to advance kinase science using The GSK Published Kinase Inhibitor Set (PKIS) (D. Drewry, Willson, and Zuercher 2014). Elkins et al. screened the 367 compound set against 224 human protein kinases at 0.1 uM and 1 uM. In the second round of PKIS, Drewy et al. predicted that 1000–1500 compounds would be required in their set to be able to selectively inhibit each of the 518 human protein kinases specifically. PKIS2 has 645 inhibitors against 392 human kinases at a dose of 1 uM. This is impressive coverage but assays at one concentration were bound to provide a poor estimate of the inhibition constant or IC50. Nevertheless their goal was different from ours; they essentially wanted to perform genetic-like experiments with drugs; we want to make use of polypharmacology to take advantage of the drugs’ properties and reduce the size of the drug ‘screen’ by at least an order of magnitude. An illustration of this difference can be found in the Gini scores of the inhibitors. A Gini score of one means the inhibitor only targets one kinase (perfect specificity); a score of zero means every kinase is inhibited equally (perfect lack of specificity!). Inclusion in the PKIS2 set required a Gini score > 0.75; the mean Gini score of our KiR3 drugs is 0.14, just enough non-specificity for a relatively small number of kinase inhibitors to cover the human kinome and just enough specificity for us using machine learning methods to deduce every kinase from the small number of experiments. We present here two new comprehensive datasets which in themselves represent a rich resource for the systems pharmacology community. The first has 257 kinase inhibitors assayed against 365 human protein kinases; the second has 58 kinase inhibitors, which were chosen to optimize our method, at ten doses (from 10 uM in a series of three-fold serial dilutions) against 369 human protein kinases. These 58 comprise what we call “Principal Compounds” - an optimal set to be used to probe the entire kinome with the reverse engineering framework defined here.

We are not alone in utilizing drugs’ properties for the purpose of deconvolving drug targets. Several studies have utilized the Anastassiadis et al. paper, for example Gautum et al. (Gautam et al. 2019) applied a machine-learning-based approach to identify cancer-selective targets in triple-negative breast cancer cell lines. Arang et al. (Arang et al. 2017) applied KiR to identify host regulators and inhibitors of liver-stage malaria and even predicted which drugs would be most effective at eliminating infection.

In order to test the power of our new set of the Principal Compounds, we applied our optimal drug set to cell motility, repeating semi-automated scratch assays we performed previously but with our updated inhibitor data now measured at multiple doses (Taranjit Singh Gujral, Peshkin, and Kirschner 2014). The scratch assay is a simple quantitative phenotype for the application of KiR and we are able to obtain reliable, repeatable dose-dependent responses. We now apply some of the latest statistical learning techniques and determine a rank-ordered list of kinases involved in cell motility. We confirm some of our previous findings and also find a few novel kinases not previously implicated in cell motility. Details of the method are shown in Figure S1.

## In vitro assays of kinase inhibitors

In order to apply regression analyses using kinase inhibitors as perturbations to the natural state of a cell, it is necessary that we know quantitatively how the inhibitors act on the largest fraction of protein kinases. It is not enough to know that a certain compound is supposed to target, say, the cyclin-dependent kinase family; our experiments will be oblivious to our preconceptions about what a target is and what is an ‘off-target’; everything is assumed to be a target. It is therefore necessary to quantitate all possible inhibitions. We will discuss some limitations of our present database in the Discussion. Therefore, it is necessary to test each compound against every one of the available kinases. To this end we collaborated with Reaction Biology Corporation (see Supplementary Methods for details) which runs high throughput *in vitro* biochemical assays of substrate phosphorylation using P33-labeled-ATP and quantifies the amount of radioactivity incorporated into substrates, which in some cases would be autophosphorylation of the kinase.

Our goal is to find a small set of inhibitors which optimally tile the space of kinase inhibition, i.e. poly-specific inhibitors with partially overlapping set of targets, where each target kinase is inhibited by multiple drugs to a different degree. Profiling at a single *ad hoc* chosen dose was meant as the first step to be followed up by a more comprehensive dose response study. To cast a wide net, initially we profiled for 257 compounds at one dose against 365 human protein kinases. Included in this set of known kinase inhibitors were approved and investigational drugs and many ‘probe’ compounds. We also included several agents with previously published kinome-wide profiling data as controls. We started with the previously published set of compounds profiled at 500 nM (Anastassiadis et al. 2011) and reasoned that compounds that did not show much activity needed to be applied at a higher dose (e.g. Go-6976), while those that were too promiscuous (e.g. staurosporine) needed to be applied at a lower dose. That gave us 10 compounds at 50 nM, 18 compounds each at 2000 nM and 5000 nM, and the remaining majority 211 compounds at 500 nM. We called this collection of inhibitors with optimal assay concentrations our “KIR2” library of drugs. Following our previously published methods (Taranjit Singh Gujral, Peshkin, and Kirschner 2014) we performed a principal component analysis to choose a subset of the highest loading variables we dubbed “PCs” (principle compounds): those compounds that capture most variation in the kinome inhibition. In addition to purely statistical considerations, we took into account biochemical properties of the drugs and supplemented several compounds we hoped to target the “dark matter of the kinome” – kinases not at all targeted by any of the original 257 in the KIR2 library. These included several non-traditional chemotypes that do not rely on classically privileged structural elements with known hinge-binding motifs. This chosen set of 58 compounds is referred to as PCs while the set of PCs at multiple concentrations as “KIR3”.

The 58 compounds of KIR3 were assayed at 10 doses across 369 human protein kinases, from 10 uM down to 1 nM in a series of three-fold serial dilutions. The activities are normalized to a control of zero drug, so we expect the points to lie in the 0–100 range. The residual kinase activity plotted against the concentration of the compound in log10 space is expected to give a sigmoid (Figure 2A). In the ideal scenario we would be able to plug in the data directly into the regression analysis. However we encountered a number of issues which we discuss below. Briefly, some drugs precipitate out of solution at higher doses, some control assays fail producing activity over 100%, some noise is injected because of numerous specific issues, such as fluctuations in the readout, plate border effects and uneven batches of protein obtained from vendors. To address these issues and arrive at robust inhibition values, we decided to fit sigmoids to the data, interpolating missing values to avoid outliers Figure 2B). Any serious deviation was grounds for rejecting the measurement. For each of the 21,000 (58 x 369) plots we fitted a sigmoid to the 10 doses plus control (Equation 1.1), where **n** is the Hill coefficient, C10 is the drug concentration in log10 space, and K10 is the IC50 in log10 space. Of the four parameters defining a sigmoid, we fixed two: the maximum and minimum were set to 100 and 0, respectively. The Hill exponent and IC50 values were fitted to the data; the permissible ranges were 0–5 and −10.5 – −3, respectively to reflect biochemically plausible ranges.

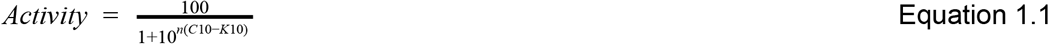

**Figure 2:**
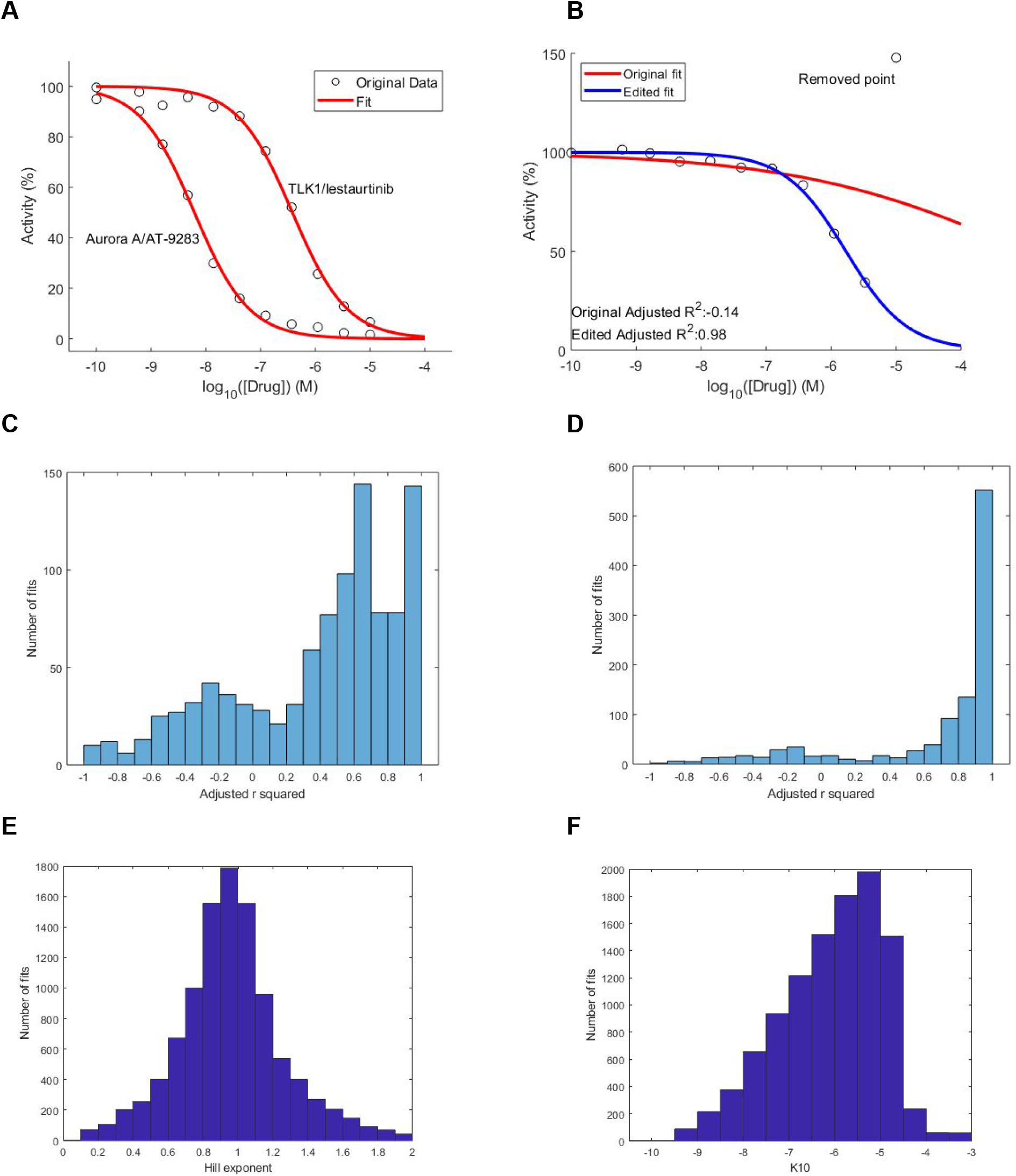
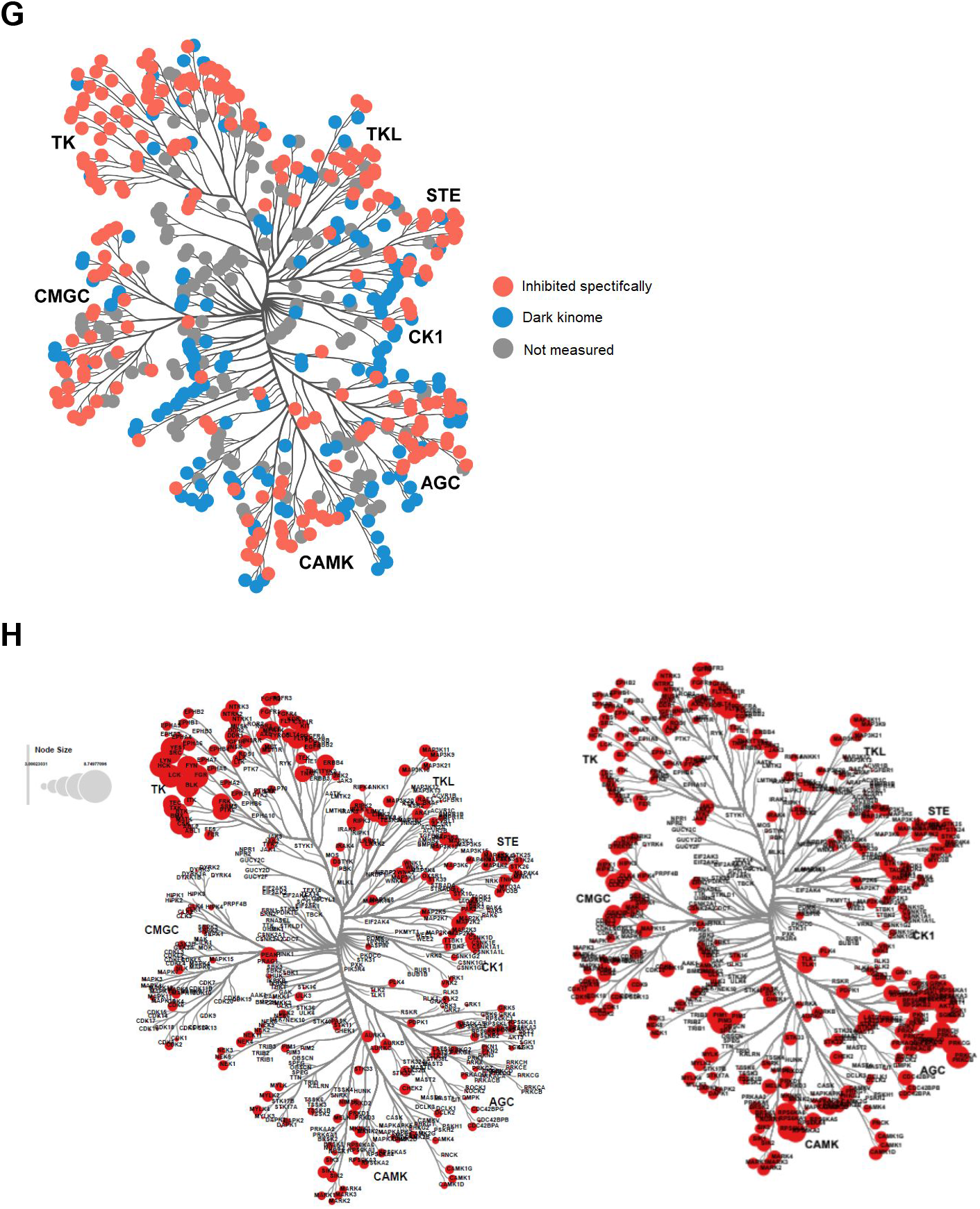
In vitro reconstitution of kinase activities for different kinase inhibitors **A)** Examples of kinase residual activity under inhibition. The activity of two kinase-inhibitor pairs illustrates the sigmoidal shape of this dependence. These particular combinations are TLK1 kinase with *lestaurtinib* and Aurora A with *AT-9283*. The activity is expressed as a percentage of control activity with zero drug. A sigmoidal fit to the data is shown. The four parameters of a sigmoid are the maximum, minimum, IC50, and Hill exponent. The maximum and minimum are set to 100 and 0, respectively; the IC50 and the Hill exponent are free parameters to be fitted; in this example they are −6.42 and 0.87 with an adjusted **r**-squared of 0.99; and −8.21, 0.87, and 0.99, respectively. The IC50 has bounds of −10.5 and −3; the bounds on the Hill exponent are 0 and 5. **B)** Example of manual curation of the data when one of the points (data points are open circles) is an outlier for the kinase RSK3 and the drug *pluripotin*. The plot is presented when the original fitted sigmoid (red) has an adjusted **r** squared < 0.7. Clicking on the point which is an outlier removes it; a sigmoid is fitted to the remaining points (blue). Removing the outlier changes the fitted IC50 by almost three orders of magnitude. **C)** Adjusted **r**-squared of fitted sigmoids before modification **D)** Adjusted **r**-squared of fitted sigmoids after modification. **E)** A histogram of the fitted Hill exponents for the data with multi-doses. The fitted values could vary from 0 to 5; they are centered on 1, which is the special case of a hyperbola. Fits where the highest dose is above 70% have not been included. **F)** A histogram of the fitted IC50 values for all of the multi-dose data. Fits where the highest dose is above 70% have not been included. **G)** Plot of kinases that we can inhibit ‘specifically’ (see text for definition). **H)** The fitted IC50 values (absolute value in log10 space) are plotted for the entire human protein kinome. The compounds are Lck inhibitor (left) and Go6850 (right).

A sigmoidal fit described most of the data (95.1 % of the 21,771 plots) well, which we determined by an adjusted **r**-squared value greater than 0.7, and needed no further modification (Figures S2D, S2E). This gave us confidence in our data and our rationale for fitting a sigmoid to each plot. For the remaining 4.9 % we manually inspected the plots and considered several modifications, described below, that could provide a better fit; after each modification we refitted a sigmoid and chose the case with the highest adjusted **r** squared value (Supplementary Table 2.1). There were numerous instances where one or two data points were clear outliers; this frequently occurred at high drug doses and could be because the drug precipitated out of solution. In these instances, we removed the outliers and refitted a sigmoid to the remaining data (Figure 2B); this accounted for 644/21771 = 2.96 % of the plots. The remaining 2 % had either normalization-type errors or ‘activation’ errors; more information can be found in the Supplemental Information, Figures S2A, S2B, and S2C and Supplementary Table 2.1. By making these modifications we improved the fitting; of all the modified plots (4.9% of the total) the mean adjusted **r**-squared value before modification was 0.06 and after the mean **r**-squared value was 0.67; the distributions are given in Figure 2C and Figure 2D. We therefore felt it was reasonable to correct the small fraction of plots that had poor fits and make these small adjustments in the data and substantially improve the data consistency.

We further can assess our fitting results by looking broadly at the free parameters that were fitted: the Hill coefficient and the IC50 values. We plotted these parameters for all kinase-inhibitor pairs, but only where the highest dose of drug gave a kinase activity below 70 % (to avoid confounding the plots with drug/kinase combinations where the drug had little effect). Plotting the Hill coefficients (Figure 2E, we see the histogram is centered around 1, which is a special case of a sigmoid, the hyperbola. In a simple biochemical binding reaction this is what we would expect. A histogram of the IC50 values is shown in Figure 2F, where we again only include drug/kinase combinations where the drug was effective. This shows that the compounds making up our optimal set have broad inhabitability of the kinome, which is desirable for our methodology. To gauge the coverage of our drugs over the human protein kinome we plot to what degree we can ‘specifically’ target a kinase (Figure 2G). For our method to give a kinase a non-zero coefficient, the kinase must both be inhibited by at least one drug/dose and that drug/dose must not inhibit too many other kinases. By this crude metric we defined a potency threshold and a specificity threshold: a kinase is inhibited when below 40 % activity, and that particular drug/dose is specific if it does not inhibit more than 20 kinases (by the same definition). We determined which kinases had at least one drug/dose that could ‘specifically’ inhibit them and overlaid this with the kinome tree (Figure 2G). We have good coverage of all of the kinase families and so our method will be suitable for a variety of cell biological problems. Of 369 kinases measured 221 (60%) could be specifically identified by this method. The rest are not inhibitable by the inhibitors and we refer to these as “dark matter of the kinome”; we know it is there, we just cannot detect it with the tools we currently have. The rest are kinases that are not assayed by Reaction Biology and include atypical kinases and often kinases of obscure function.

The coverage of the inhibitors is further exemplified by overlaying the fitted IC50s onto the human protein kinome (see Figure 2H for two examples). The absolute values of the fitted IC50 values in log10 space are plotted on the human protein kinome tree. As an example of the type of differential coverage we find for single inhibitors, one can see the LCK inhibitor mainly inhibits the subfamily containing LCK and also the TK family of kinases, whereas the Go6850 inhibitor is less specific to one family but does target the AGC and CAMK families preferentially. It is these features of the inhibitors in our optimized set that makes them ideal for KiR.

Additionally to judge the quality of the measurements, the assay was repeated for all of the compounds and 50 of the protein kinases. There we have two sigmoids for a given kinase-inhibitor pair, the fit with the largest adjusted **r**-squared was chosen for future analyses; of 2950 repeated kinase/drug combinations, 1433 (48.6 %) were chosen over the original data. Comparing the repeats shows there is good agreement; when we plot the IC50 values for the original and repeated data against each other the points predominantly fall on the diagonal (Figure S2F). 75.6 % of the comparisons fall within half an order of magnitude of each other. Where the points do not fall on the diagonal, in many cases it is because one point is at the upper extreme of the fitting parameter range (which is where the points remain at 100 %). This suggests the disagreement is because the assay completely “didn’t work”, perhaps because the drug precipitated out of solution, rather than there being a quantitative difference in results. Indeed, when we only consider points where both original and repeat have IC50s lower than −4.5 in log10 space, 94 % of the IC50s fall within half an order of magnitude of each other. The problem of differences between repeated data is therefore mitigated by choosing the fitted sigmoid between the repeats which gives the highest adjusted **r**-squared value. A further way to compare ‘repeats’ of the data is to compare the original assay, where we assessed 257 compounds at one dose, with the assay for a subset of those compounds at several doses. We took the dose used in the original assay and extracted the degree of inhibition from the fitted sigmoid plot at the corresponding doses for the 365 kinases and 50 drugs shared between these datasets; this also gives good agreement (Figure S2G). We expect this comparison to be noisier than the above IC50s comparison as the ‘original’ here is only one datapoint rather than a value from 11 data points. We observe that 95.1 % of comparisons are within 20 % of each other. This additional comparison further reassures us of the quality of our dataset and parameter fitting.

In summary, we set out to improve the set of kinase inhibitors used in KiR to give broad coverage of the human protein kinome. That said the kinases that have been best covered are ones of the widest distribution and the most sensitive function. We to some degree already achieved much of this goal. The dose–response curves give us much greater confidence compared with using single-dose values, and we were able to fit sigmoids to these data (with some modifications). By assessing the fitted IC50s and overlaying them with the kinome tree we can obtain the present state of coverage. We therefore are confident to begin to use the KIR3 set of 58 kinase inhibitors up to 10 uM as a discovery toolkit.

## Using our optimal drug set to perturb a quantitative phenotype: cell migration in a wound healing assay

Cell migration is a complex process that engages many signaling pathways within a cell, from regulation of the cytoskeleton to G-protein signaling to cell polarity. It is known that many protein kinases play a role in this regulation (Ridley 2003). Cell migration is often studied using the so-called “wound healing” or “scratch assay”. In the scratch assay, a uniform region of a confluent cell monolayer is cleared of cells via a reproducible scratch device (Figure 3A and Supplemental Information). Reoccupation of the cleared scratch area occurs over time as cells fill in this blank region through migration and this process can be quantitated using simple forms of image analysis. The process of “wound healing” can be affected by gene knock-downs and drugs. In a previous study, we searched for kinases involved in migration by applying KiR to scratch assay data using a significantly smaller kinase inhibitor dataset (Taranjit Singh Gujral, Peshkin, and Kirschner 2014). Here, we revisit the role of kinase activity in cell migration by application of KiR3 using our new optimal set of kinase inhibitors at several doses.

**Figure 3:**
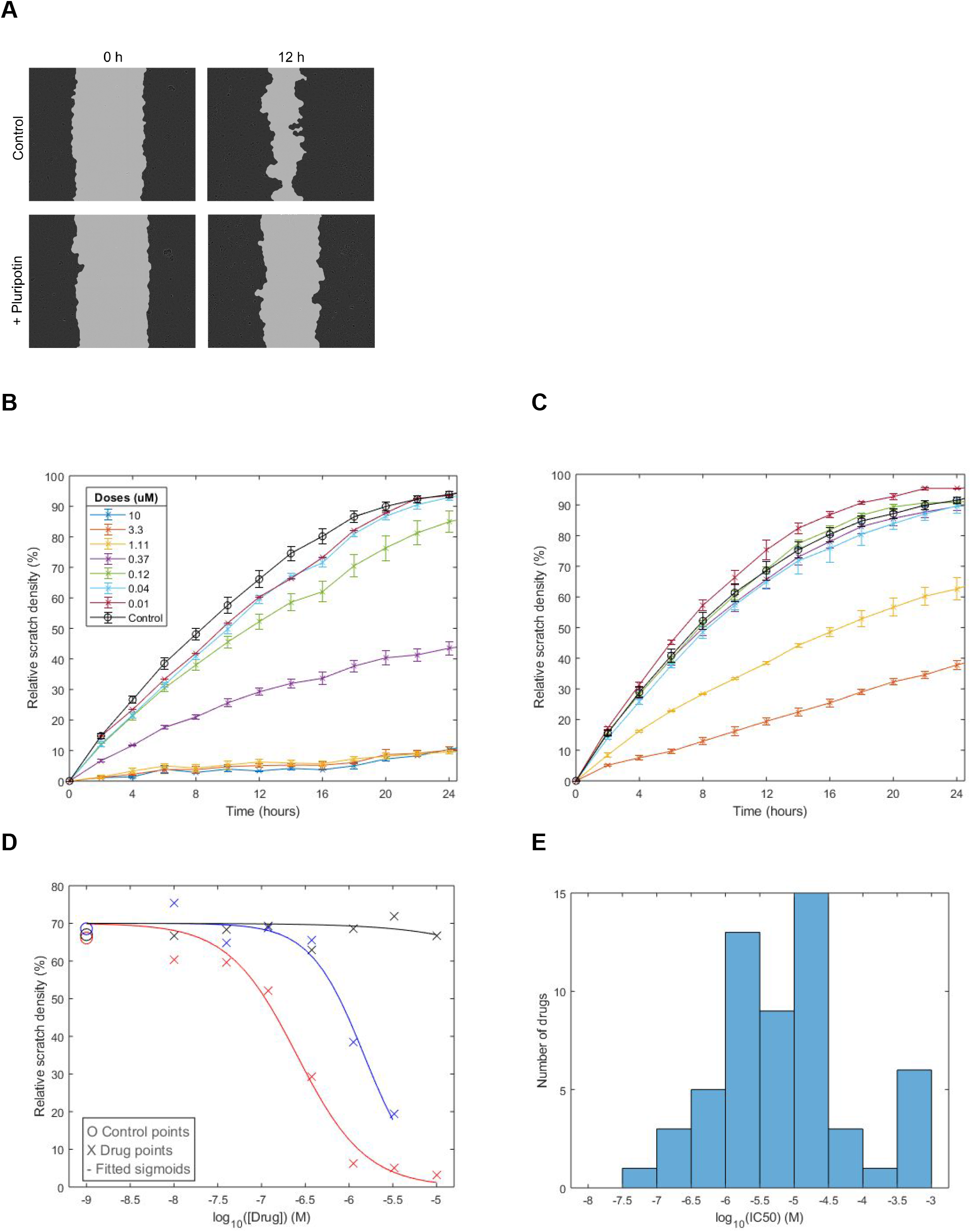
**A)** Example images of one control and one inhibitor-treated (370 nM *pluripotin*) well at two time points. The segmentation of the image is shown with the cellular region (outside of the scratch), dark gray and the acellular scratch region is shown in light gray. The initial scratch is made (left) and is uniform for all wells (left images). After 12 hours, the control well (top right) has around 70 % relative scratch density (see Supplemental Information for definition). In the presence of *pluripotin*, the scratch closure at 12 hours (bottom right) is impaired compared with control (top right). **B)** Example time course of the effects of *pluripotin* and **C)** *nintedanib* on cell migration for seven doses plus control (DMSO). The highest dose of *nintedanib* was toxic and therefore removed from the analysis. The highest dose of *pluripotin* is 10 uM and a three-fold serial dilution was performed. The scratch was made and the compounds added just prior to time point zero. Images were taken every two hours and the relative scratch density calculated as described in the Supplemental Information. Doses are in triplicate; control has twelve wells across four drugs per plate. Means and standard errors of three replicates (12 for control) are plotted. **D)** Representative examples of dose-response data revealing a range of effects of kinase inhibitors on cell migration. A plot of the relative scratch density for *pluripotin* (red), *nintedanib* (blue) and *apatinib* (black) with the drug dose on the x-axis. A single data point is required at each dose for the regression analyses; the time point with maximum variation between doses was found to be when the control plot was closest to 70 %. A sigmoid was fitted to each of these plots (56 – one for each drug). Of the four free parameters for the sigmoid, two – the maximum and minimum – were fixed at 70 and 0, respectively; the IC50 and Hill exponent were fitted and had ranges of −10.5 to −3 and 0 to 5, respectively. **E)** The distribution of the IC50s obtained for all of the inhibitors used in the scratch assay.

The scratch assay used here has several features that make it especially attractive for use with KiR (see Supplement for experimental details). It can be performed in 6 96-well plates per run, achieving a throughput sufficient for testing multiple inhibitors in a dose-response format with replicates. The multiwell format requires small volumes, minimizing consumption of expensive compounds. Scratches can be made reproducibly both across plates and from run to run allowing for comparison of measurements made from different plates or even on different days. Finally, sensitive quantitation allows for the detection of small differences in cell migration that result from treatment with inhibitors. In this experiment, seven doses were tested in triplicate for each 56 inhibitors in our optimal drug set and along with DMSO controls using the Hs578t cell line. This cell line, derived from mammary carcinoma, was also used in our previous work (Taranjit Singh Gujral, Peshkin, and Kirschner 2014).

As a quantitative metric of cell migration we computed the relative scratch density, which is roughly the ratio between the cell density within the region inside and the region outside the original scratch (see Figure 3A and Supplemental Information). Treatment with kinase inhibitors may reduce proliferation or even lead to cell death. Although we did see some clear signs of inhibitor toxicity, it only occured at high concentrations of select inhibitors and these concentrations were not used in the analysis (we removed 38 such data points). The relative scratch density, which, to a degree, accounts for changes in confluency outside of the scratch attempts to correct for inhibitor toxicity in the cases where it may not be obvious and therefore manually removed.

We can assess the relative scratch density over time for the different drug doses (Figures 3B, 3C, and S3D). In control conditions, scratch closure begins within the first hours and the rate of closure is maximal for the next 8–12 hours. With increasing time, the closure rate begins to slow and closure is complete within 36 hours. These kinetics are altered by the presence of inhibitors but in ways that depend upon the concentration. We define the regression phenotype for a given concentration of inhibitor to be the percent scratch density at the time point where the scratch density of the control is equal to 70%. This value was chosen as it reveals small, dose-dependent changes in scratch closure that can be obscured at later time points, especially in the presence of low concentrations of inhibitors (Figure 3B). This definition results in a sigmoid with a maximum of 70%, a minimum of 0%, and clear, reproducible dose-response behavior (an example of such a plot is shown in Figure 3D).

Kinase inhibitors produced a range of effects on scratch closure which were dose-dependent and well-described by a four-parameter sigmoid curve fit (see Supplementary Information). Fitting produced IC50 values that fall in a reasonable range of mid-nM to low-mM with the majority in the low μM as shown in Figure 3E. Thirty-one of 56 inhibitors tested had fitted IC50 values less than the highest dose of inhibitor applied, indicating that more than half of the inhibitors produced strong effects on cell migration and could be reliably quantitated. Although it is reassuring to see a large number of inhibitors producing strong effects, it should be noted that all inhibitors used in KiR are poly-specific; inhibitors that show no effect provide very important information regarding kinases that have no effect on migration. We conclude that the scratch assay is well behaved and capable of providing high quality information for use in identifying kinases responsible for the phenotype.

## Identification of kinases with major roles in cell migration: use of Elastic Net Regularization

With the matrix of kinase inhibitor specificity and the scratch assay phenotype data at hand, we can apply statistical learning methods to reverse engineer the kinases affecting the phenotype. We have information on how each inhibitor affects the purified kinases in our database and information on how each of our PC inhibitors affects the specific phenotype. From these two we wish to model how individual kinases affect the phenotype (Figure 4A). Previously, we used a linear model with penalty terms for the number of coefficients included (Gujral2014). Each datapoint, **y**_i_, is modelled as the sum of the activities of each kinase multiplied by a fitted coefficient. We transform the scratch assay data using the logit function; the data points are then between 0 and 1. The objective is to minimise the sum of squares of the difference of the data, penalized for model complexity (regularization) (Equation 4.1). Alpha is a weighting for the type of penalty used in the model; 0 is ridge regression and 1 is lasso regression. Lambda is a complexity parameter to determine the tradeoff between the number of variables chosen and the goodness of fit of the model, n is the number of data points, and p is the of variables (kinases).

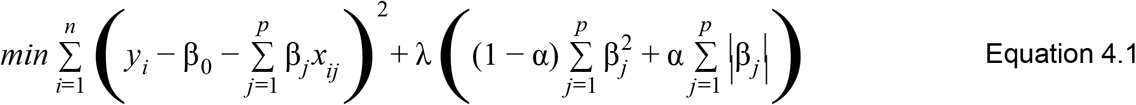

**Figure 4:**
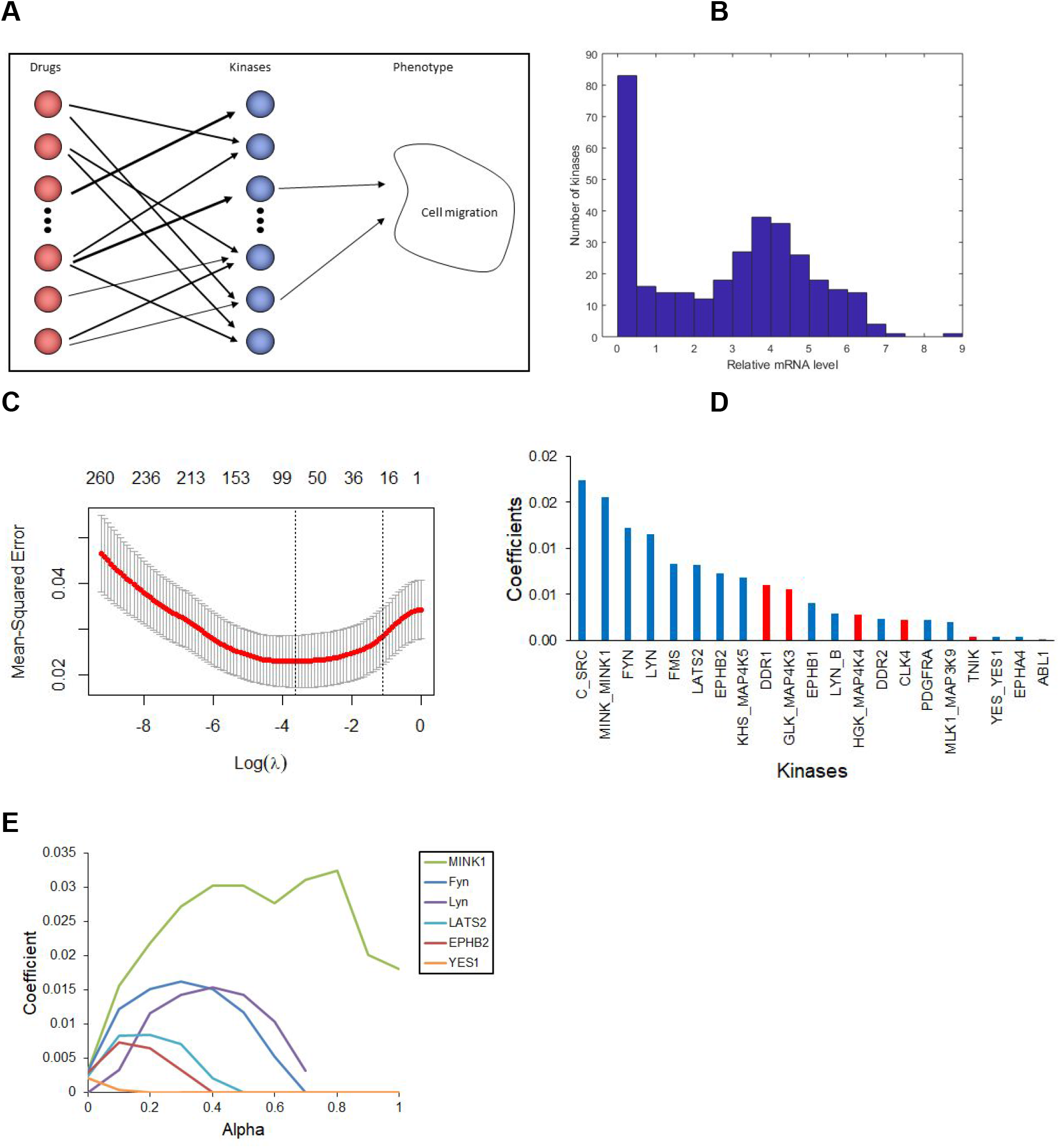
Elastic net regularization using the kinase activity data together with quantitative cell migration data. **A)** schematic of the rationale behind using kinase inhibitors to identify kinases involved in a particular phenotype. The inhibitors target kinases in a known way, and they also affect the phenotype of interest in a measurable, quantifiable manner. The mapping between phenotype and kinases can then be determined; there are many methods available to do this – we analyze several here. Boldness of arrows depicts differing magnitudes of effect. **B)** The expression of mRNA in Hs578t cells. **C)** Result of leave-one-out cross-validation using glmSparseNet. The mean-squared error is plotted against the log of the regularization parameter lambda. The value of lambda is chosen so that it is within 1SE of that which minimizes the mean-squared error and the coefficients at this value are determined. Alpha = 0.1. **D)** A ranked list of kinases obtained in our analysis. Blue: kinases have a bona fide role in cell migration or were found in our previous analysis. Red: no known role in cell migration. Other kinase coefficients are equal to 0. The majority of the kinases we found either have *bona fide* roles in cell migration or were found in our previous analyses. **E)** Kinase coefficients at lambda_1se for different values of the elastic net parameter alpha. As alpha tends towards one many of the coefficients become zero, a feature of LASSO regression.

Before running the regularized regression analyses one important step should occur: to consider which kinases in our matrix are actually expressed in the cells we are using. In our previous implementation of KiR we used mRNA expression as a proxy for protein expression; we do the same here but we assessed the kinases in our matrix using data from HMS LINCS (HMS dataset ID 20348), where the counts were normalized to reads per kilobase of transcripts per million mapped reads. In this single dataset we found expression data for 366/369 of the protein kinases in our matrix (figure 4B). Analyzing the histogram, we see a bimodal distribution and therefore decided to choose 0.5 units as a cut-off, which led to 83 kinases being removed from the matrix (the three without mRNA data were preserved in our matrix). The remaining 286 kinases were the ones we used in all subsequent analyses.

Regularized regression methods differ in how they handle dependent variables. E.g. a pair of orthologous kinases could be closely related in how they react to the same drugs, thus one of the kinases possibly substituting another with its explanatory power. Depending on the regularization penalty only one or both in such a pair would be included in the list of selected regressors. Moreover kinases weakly correlated with one another in their response create a web of relationships with it’s “hubs” and terminals, where a hub is one kinase correlated with a few others stronger than each of these are with one another. Since we are interested in identifying the kinases that are likely “first responders” early in signaling cascades, we used the *glmSparseNet* package in R with leave-one-out cross-validation (LOOCV); this package is advantageous over *glmnet* which we used previously as it takes better account of correlated variables (Veríssimo et al. 2018). Since the data points from the same drug are not independent of each other, we removed all points from the same drug in the training set during LOOCV. The mean-squared error for different values of the penalty parameter, lambda, is shown in Figure 4C for alpha = 0.1. The coefficients of the kinases from this model are shown in Figure 4D. We explored the full range of the elastic net parameter *alpha* and the resulting kinase coefficients. The six examples in Figure 4E show that some kinases are more robustly predicted; MINK1 has non-zero coefficients for all values of alpha, whereas YES1 is non-zero for only the lowest two values of alpha.

In the list of kinases with non-zero coefficients we find kinases with *well-documented* roles in cell migration. In our previous application of KiR, we identified the SRC family member Fyn as the kinase in a non-canonical Wnt signaling pathway, phosphorylating Stat3 (Taranjit S. Gujral et al. 2014; Taranjit Singh Gujral, Peshkin, and Kirschner 2014). We again find this kinase, along with its family members LYN, LYN B, SRC, and YES1. The Ephrin receptors are also known to play a role in cell migration (Park, Inji and Lee, Hyun-Shik 2015); there are three in our list of predicted kinases (EPHB2, EPHB1, and EPHA4). PDGFRA is also known to regulate cell migration (Yu, Moon, and Kim 2001)

Importantly, our expanded matrix that includes additional kinases allowed discovery of four new kinases that were not in the previous matrix: LATS2, DDR1, GLK/MAP4K3, and TNIK. LATS2 kinase is also known to play a role in cell migration, potentially through the Hippo pathway, although it is thought to act to inhibit cell migration (Visser and Yang 2010). DDR1 has also been shown to play a role in cell migration, promoting migration by activating the RhoA/ROCK/MAPK/ERK pathway (Azreq et al. 2016); we found its closely related family member DDR2 now and previously. GLK/MAP4K3 was found to promote lung cancer metastasis by phosphorylating and activating IQGAP1 (Chuang et al. 2019). TNIK is an activator of Wnt target genes and therefore also promotes cell migration (Mahmoudi et al. 2009). This demonstrates an improvement on our previous method. A curious kinase in our list is CLK4, which, intriguingly, is involved in the regulation of spliceosome formation (Schultz et al. 2001); it is unclear if it also plays a direct role in cell motility.

## Model validation

In addition to comparing our results with the literature and our previous results, we sought to validate our findings by repeating the scratch assay with siRNA for six of the kinases in our list (Figure 5). Cells were seeded and treated with siRNA for 2–2.5 days. Knockdown of the SRC family members Fyn, Lyn, and YES1 each resulted in slower cell migration, as did knockdown of the ephrin EPHB2. MINK1 reduced migration, as we found previously. YES1 kinase knockdown did not slow cell migration; it actually increased the rate of the scratch closure at later time points. This is not completely surprising: our model did not robustly predict YES1, and it is known to have an inhibitory role in cell migration (Visser and Yang 2010).

**Figure 5:**
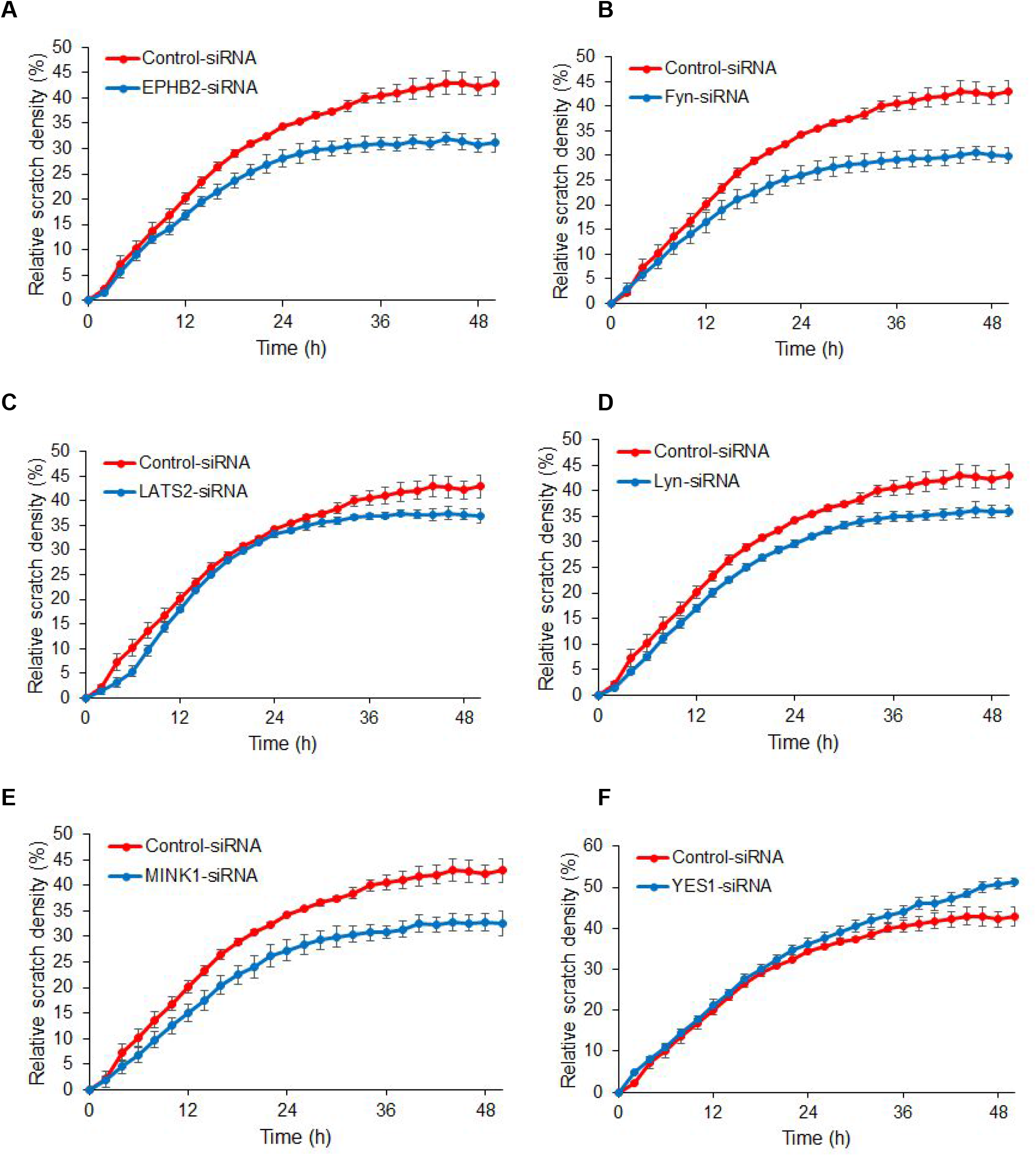
Scratch assay with siRNA of selected kinases.

We were able to use our curated matrix of kinase activities with our optimal drug set together with our quantitative phenotypic values from the scratch assay using the same optimal drug set to perform regression against the kinome. We used an updated statistical learning package, glmSparseNet, and explored the range of penalty terms to determine a list of kinases, together with their coefficients, involved in cell migration.

Our goal was to improve upon our initial successful attempt at using polypharmacology to deconvolve protein kinases involved in cell migration. Pharmacology has many very specific advantages and disadvantages over other screening methods such as genetics: Drugs that inhibit kinases can target post-translational modification as well as translational control and protein degradation. Previously, we utilized a publicly available kinase inhibitor dataset to regress quantitative phenotype values against kinase activity levels. We sought to construct a better dataset for our method. We first made in vitro measurements of 257 protein kinase inhibitors at one dose against more than 70% of the human protein kinome. We chose a subset of those inhibitors and assayed them further: at ten doses across three orders of magnitude. We have good coverage across all kinase families and the dose-response relationships of the kinases are as we expect.

To determine the functionality of our purportedly improved dataset, we repeated the scratch assay that we employed previously with all drugs in our set at multiple doses. These experiments gave dose-dependent responses of the cellular phenotype and more than half of our drug set had some effect on cell migration. We applied KiR, using glmSparseNet, which is an improved statistical learning method over glmnet which we used before. The output of KiR is a rank-ordered list of kinases. A major finding from our previous application was the SRC family member, Fyn – it is part of a non-canonical Wnt signaling pathway and phosphorylates Stat3. We now see Fyn in the top three kinases, which may be an indication of how improved precision in characterizing the panel of inhibitors against a large fraction of the kinome improves the identification of what is now a highly validated component of an important Wnt pathway that plays a role in metastasis. Previously Fyn was ranked 13th with a coefficient an order of magnitude smaller than the largest one; it now ranks third and is accompanied by close members of the same family. In retrospect it is surprising we were able to see it at all with the early data set..

We envisage many applications of this rich dataset, from more cellular phenotypes to predicting drug behaviour based on structure. In addition, we are working on an approach that does not have such a strict workflow of doing all experiments before any analysis. Depending on the question the biologist asks, we can optimize an initial small set of drugs that should be used first; for example we can optimize a drug set to answer the question ‘which kinase family is responsible for a phenotype?’

## Acknowledgements

This work was supported by the intramural programs of the Center for Cancer Research, National Cancer Institute and the Division of Preclinical Innovation, National Center for Advancing Translational Sciences of that National Institutes of Health.

## Supplemental Information

### RBC assay details

The in vitro reconstitution assays to determine the efficacy of kinase inhibitors on human protein kinases were performed by Reaction Biology Company using their HotSpot Kinase Assay Protocol. The Base Reaction buffer consists of: 20 mM Hepes (pH 7.5), 10 mM MgCl2, 1 mM EGTA, 0.02% Brij35, 0.02 mg/ml BSA, 0.1 mM Na3VO4, 2 mM DTT, 1% DMSO. Required cofactors were added individually to each kinase reaction. Testing compounds were dissolved in 100% DMSO to specific concentration. The serial dilution was conducted by epMotion 5070 in DMSO.

The reaction procedure was:

1. Prepare substrate in freshly prepared Reaction Buffer
2. Deliver any required cofactors to the substrate solution above
3. Deliver kinase into the substrate solution and gently mix
4. Deliver compounds in 100% DMSO into the kinase reaction mixture by Acoustic technology (Echo550; nanoliter range), incubate for 20 min at room temp
5. Deliver 33P-ATP (Specific activity 10 Ci/l) into the reaction mixture to initiate the reaction
6. Incubate for 2 hours at room temperature
7. Spot the small portion of the reaction onto P-81 ion-exchange filter paper (Whatman)
8. Wash out the unbound phosphate on the filter paper in 0.75% phosphoric acid buffer for 3 times and dry
9. The radioactivity remained on the filter paper was measured
10. Kinase activity data were expressed as the percent remaining kinase activity in test samples compared to vehicle (dimethyl sulfoxide) reactions. IC50 values and curve fits were obtained using Prism (GraphPad Software).

### Modifying data to fit sigmoids for the assay of kinase inhibitors using an in vitro reconstitution

For each plot we considered a normalization-type error. The mean of the three lowest doses (not including control) was calculated; if this was below 90 % or above 110 % and all three points were within ten units of each other then we considered these to have a normalization error. We calculated a multiplying factor so that the mean of the three points was 100 and then applied this to all of the data points. We did a similar normalisation at high doses. If the points corresponding to the three highest doses had a mean greater than 10 or below −10 and were within 10 percentage points of each other, a multiplication factor was applied to every point which brings the mean of these three points to 0. A further kind of ‘error’ is activation; in some cases, at intermediate doses the activity of the kinase was above 100 % compared with control. This is problematic for fitting a sigmoid which is constrained to be between 100 and 0; as such any point above 150 % was set to 100 %.

### Scratch assay

All scratch assays were done using an IncuCyte FLR (Essen BioScience) and it’s associated Cell Migration kit. This system allows for a throughput of six 96 well plates per run, with each well imaged every two hours. Cells are plated at confluency in 96-well IncuCyte ImageLock plates and allowed to settle and form monolayers overnight at tissue culture conditions. The next morning, scratches are introduced in parallel using the IncuCyte WoundMaker, which produces scratches that are regular in size and location within the well. The medium is then aspirated to remove unattached cells and debris and replaced with media containing serially diluted kinase inhibitors. The plates are then immediately placed within the IncuCyte and imaging initiated.

The IncuCyte Cell Migration kit provides an analysis module that automates the analysis of images of cell migration into the scratch. Each image is segmented into cellular regions and the scratch, or areas without cells. From the segmented images, the module produces three independent metrics for each well at each time point: the scratch width, the confluence of cells within the scratch, and the cellular confluence outside of the scratch. The relative scratch density is calculated from Equation 3.1:

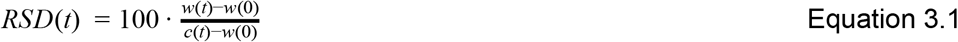

Where w(t) is the density of the scratch region at time t (the confluence of cells within the scratch), and c(t) is the density of the region outside of the scratch at time t. The variable w(0) here is simply the density of the scratch at time zero and is typically quite small. It is included to correct for potential heterogeneity in removal of cells by the scratch. The relative wound density is therefore a measure of the cell density in the wound compared with the cell density outside of the wound. Although this definition lacks an explicit measure of cell proliferation or death, it provides some measure of compensation for changes in birth or death rate in the presence of kinase inhibitors. See for example, the highest concentrations shown in Figure S3A where the confluence of cells outside of the original scratch decreases with time. We then further process the relative scratch density metric by identifying the time point at which the relative scratch density of the associated control is equal to 70% and reading off the RSD of the inhibitor-treated wells at this time point.

### Fitting sigmoids to the relative scratch density dose-response data

A sigmoid was fitted to the dose-response data for each inhibitor. The sigmoid model used is the standard four parameter model. Two of the four parameters of the model, the maximum and minimum responses, were fixed to 70% and 0%, respectively, according to the constraints arising from our definition of the response. These constraints leave two free parameters remaining, the so-called Hill exponent and the IC50, to be fitted from eight data points. The fitting was performed in Matlab, using the built in function ‘fit.m’ and the two free parameters were constrained with ranges of 0 to 5 for the Hill coefficient and −10.5 to −3 M for IC50. The constraining of the high end of the IC50 to 1mM results in essentially assigning a value of 1mM to any inhibitor that does not produce an effect (i.e., has a flat dose-response curve). For curves with fitted IC50 values below the maximum dose of drug used (10 uM), the mean adjusted **r**-squared value was 0.72.

### Elastic net regularization

Kinase coefficients obtained using glmSparseNet at lambda_1se with alpha=0.1, network=sparsebn.

**Supplementary Table 2.1:** Overview of the modifications made to fit sigmoids to the data.

| Modification | Original | Repeat | Total |
| --- | --- | --- | --- |
| Rescaling; mean of three points at low inhibitor values below 90 %. | 190 | 36 | 226 |
| Rescaling; mean of three points at low inhibitor values above 110 %. | 48 | 15 | 63 |
| Rescaling; mean of three points at high inhibitor values below -10 %. | 0 | 0 | 0 |
| Rescaling; mean of three points at high inhibitor values below -10 %. | 31 | 1 | 32 |
| Obvious outlier points manually removed. | 575 | 69 | 644 |
| Activation removed; points above 150 % set to 100 %. | 80 | 21 | 101 |
| No modifications | 19414 | 1291 | 20705 |
| **Total** | **20338** | **1433 (2950 total)** | **21771** |

**Figure S1.**
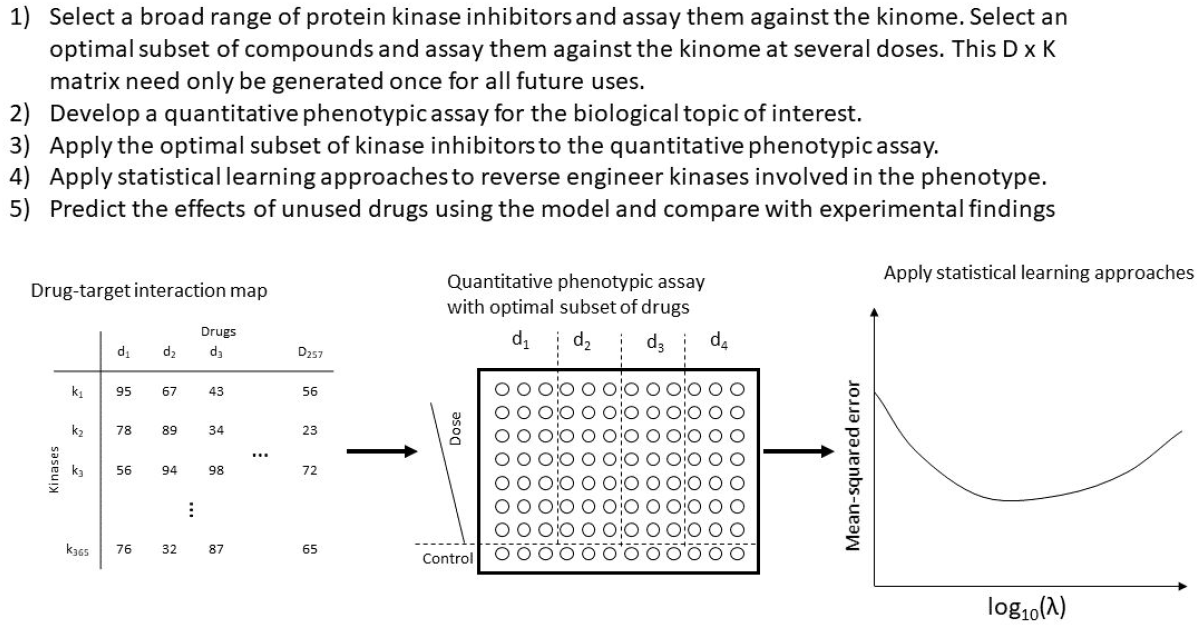
A step-by-step method of exploiting polypharmacology for drug–target deconvolution.

**Figure S2:**
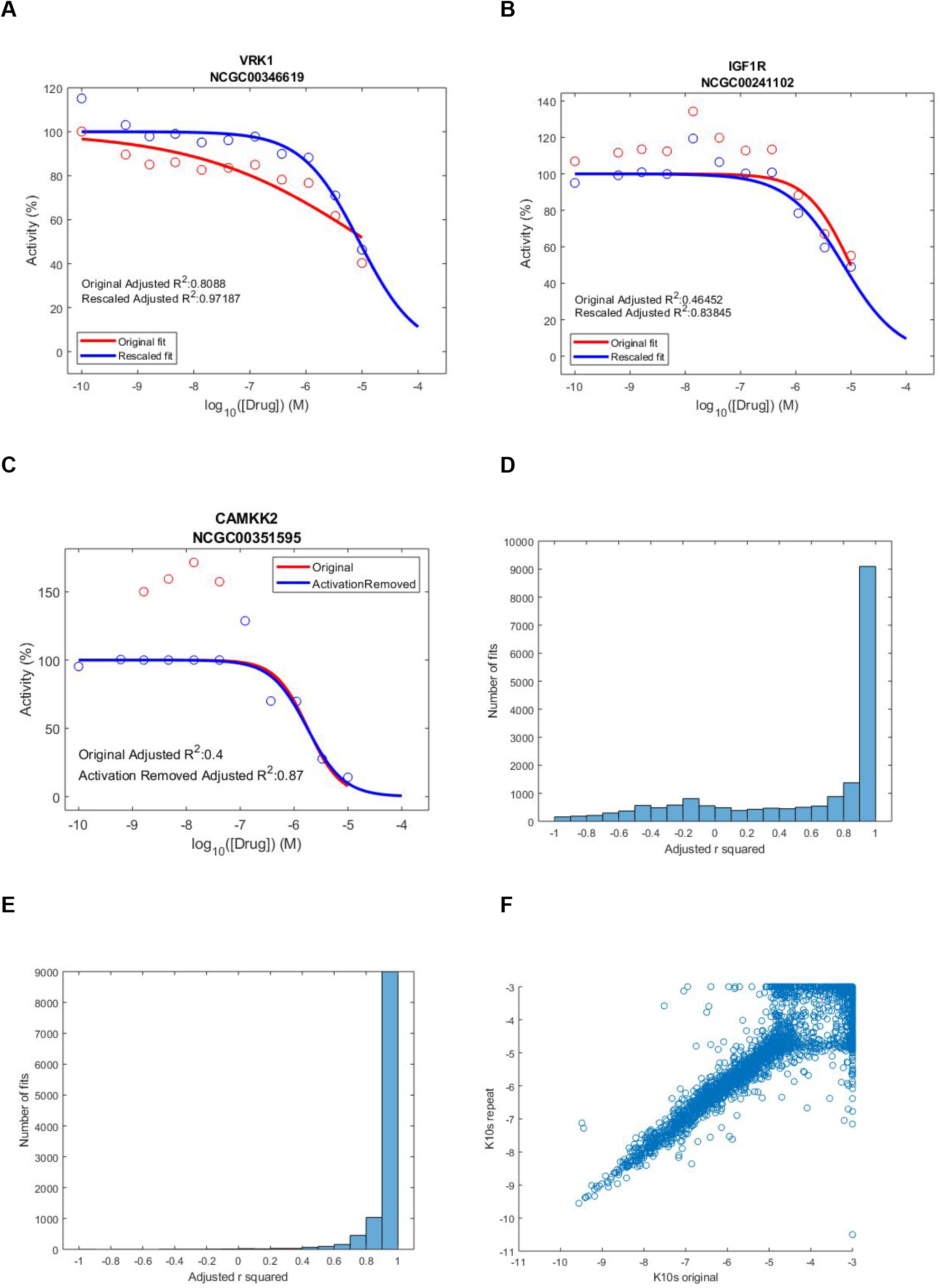
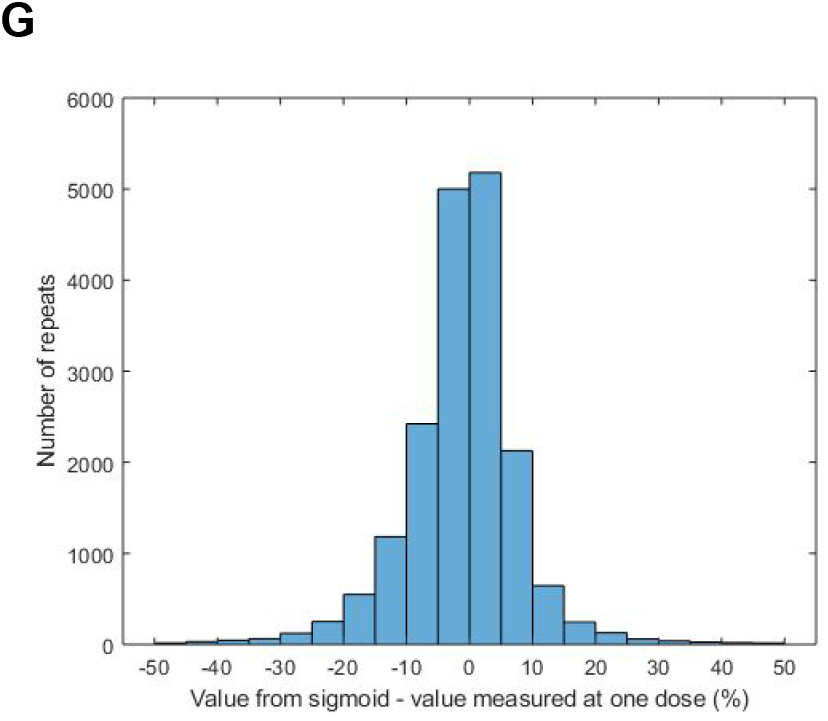
Modification of some of the kinase activity dose-response plots to improve sigmoid fitting. **A)** For each plot an automated check was performed for the first three doses above control. If their mean was below 90 % (top left) and they were within ten percentage points of each other, a multiplication factor was determined so that the mean of those three points was 100 %. **B)** Similar to (A) except the points were rescaled if the mean of the three points was above 110 % and they were within ten percentage points of each other. **C)** Removal of ‘activation’. If any value was above 150 %, it was set to 100 %. **D)** Histogram of the adjusted **r**-squared values for fitting of all of the unmodified plots. **E)** Histogram of the adjusted **r**-squared values for fitting of all of the unmodified plots and with the exclusion of plots where the highest dose resulted in greater than 75 % activity at the highest dose reading from the fitted sigmoid. **F)** Comparison of the fitted IC50 values (in log10 space) comprising the original and repeated data. **G)** Comparison of the activities in the first run of the assay with the same dose read-off from the fitted curve of the second run of the assay. The value for the one dose was subtracted from the respective value read off from the fitted sigmoid. Differences for the shared 50 drugs by 365 kinases are plotted.

**Figure S3:**
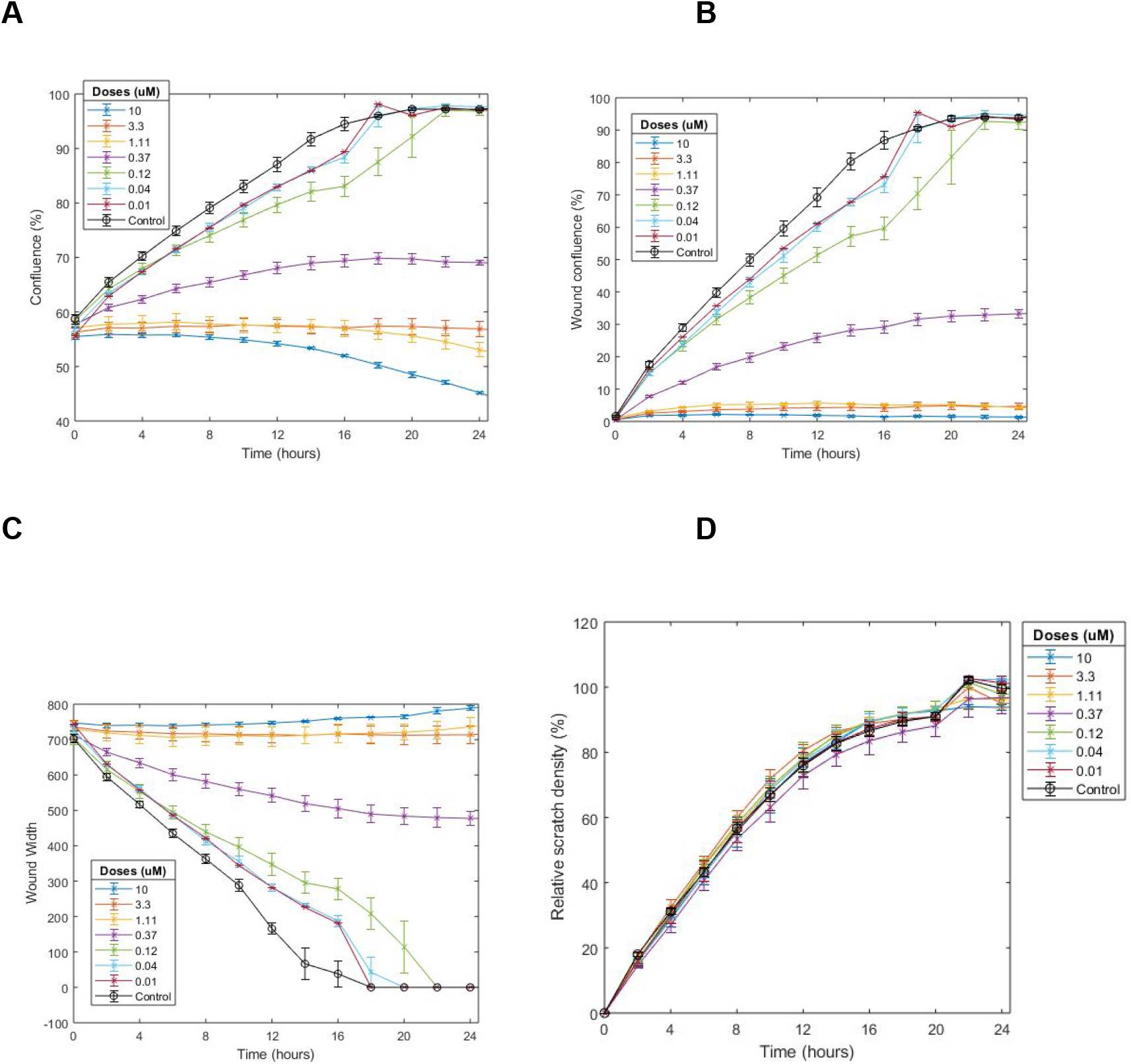
Scratch assay time courses for different phenotypes. **A), B), and C):** Example plots of other phenotypes (confluence, wound confluence, and wound width) associated with the IncuCyte scratch assay for the kinase inhibitor *pluripotin*. **D)** Time course of relative scratch density for *apatinib* (values at t = 12 hours are plotted on Figure 3D, black).

